# Mechanics and Differential Healing Outcomes of Small and Large Defect Injuries of the Tendon-Bone Attachment in the Rat Rotator Cuff

**DOI:** 10.1101/2020.07.03.184499

**Authors:** Anna Lia Sullivan, Ryan C. Locke, Rachel K. Klink, Connor C. Leek, Megan L. Killian

**Affiliations:** Department of Biomedical Engineering, University of Delaware, Newark, Delaware 19716; Department of Physics and Engineering, Taylor University, Upland, Indiana 46989; Department of Orthopaedic Surgery, Michigan Medicine, Ann Arbor, Michigan 48109

**Author notes:** CORRESPONDING AUTHOR: Megan L. Killian, PhD, University of Michigan Medical School, Department of Orthopaedic Surgery, 2021 BSRB, 109 Zina Pitcher Place, Ann Arbor, Michigan 48109. AUTHOR CONTRIBUTIONS: ALS, RCL, RK, and MLK designed the experiments. ALS, RCL, and RK acquired the data. All authors analyzed and interpreted the data. ALS drafted the manuscript. RCL, RK, CCL, and MLK edited the manuscript. All authors have read and approved the final submitted manuscript.

**Keywords:** tendon, healing, attachment, biomechanics, animal model

## Abstract

The size of rotator cuff tears affects clinical outcomes following rotator cuff repair and is correlated with risk of re-injury. This study aimed to understand how defect size influences the structural and mechanical outcomes of the injured rotator cuff attachment *in vivo*. We used our previously established model of full-thickness injury of the rotator cuff tendon-bone attachment in Long Evans rats to compare differences in healing outcomes between small and large defects. Biomechanical properties, gross morphology, bone remodeling, and cell and tissue morphology were assessed at 3- or 8-weeks of healing. At the time of injury (no healing), large defects had decreased mechanical properties compared to small defects, and both defect sizes had decreased mechanical properties compared to intact attachments. The mechanical properties of the defect groups were comparable after 8-weeks of healing and significantly improved compared to no healing but failed to return to intact levels. Local bone volume at the defect site was higher in large compared to small defects on average and increased from 3- to 8-weeks. Contrastingly, bone quality, measured as bone volume percentage and trabecular morphometry, of the total epiphysis and greater tubercle decreased from 3- to 8-weeks of healing and these changes were not dependent on defect size. Qualitatively, we observed that large defects had increased disorganized collagen and neovascularization compared to small defects. In this study, we demonstrated that not only small but also large defects do not regenerate the mechanical and structural integrity of the intact rat rotator cuff attachment following healing *in vivo*.

**Statement of Clinical Significance:** Our rat model of full-thickness rotator cuff tears may be beneficial to understand and prevent tear enlargement *in vivo*.

## INTRODUCTION

Small rotator cuff tears are one of the most common shoulder injuries.^1^ These tears can limit normal function of the shoulder^2^ and propagate into larger tears that require surgical repair.^3–6^ The size of the initial tear prior to repair plays a critical role in post-operative healing and pain scores, as larger tears have been correlated with decreased tendon integrity, decreased range of motion, and an increased rate of shoulder osteoarthritis progression.^3,7,8^ Small tears are often treated conservatively to eliminate pain. The outcomes of conservative treatments are highly variable,^4,9^ and there is a high risk of tear propagation.^10^ Larger tears commonly require surgical repair for functional reattachment of the injured tendon to bone.^11–15^ Little is known about critical size of full-thickness tears in a single tendon that propagate into larger, more problematic tears *in vivo*. This study aimed to understand how defect size differentially affects healing of the attachment between tendon and bone in an *in vivo* rotator cuff injury model of full-thickness tears.

The rat is a commonly used animal model to study rotator cuff tears due to anatomical similarities to the human rotator cuff.^15^ Previous studies using cadavers or animal models to study rotator cuff tears have shown that strain concentrations at the tendon-bone attachment are elevated with increasing tear size.^16–20^ The majority of animal models used to study rotator cuff healing rely on complete tendon tears with sutured reattachment of tendon to its bony footprint to mimic the clinical scenario of rotator cuff repair.^21–31^ Yet, few studies have investigated non-operative healing of small rotator cuff injuries *in vivo*.^18,19,32^ While simulating the clinical scenario of rotator cuff tear and repair has benefits for understanding healing mechanisms of surgical repair, several unanswered questions remain regarding the regenerative capacity of the native tendon-bone attachment without surgical repair. We have previously observed that full-thickness injuries of the tendon-bone attachment heal via fibrosis and result in significant reductions in bone quality.^19^ Additionally, we have shown that non-healed small defects at the tendon-bone attachment lead to decreased strength and the formation of strain concentrations across the remaining intact attachment.^18^ However, our previous studies only explored outcomes associated with healing and mechanical properties using a small defect and also did not compare mechanical outcomes following healing without repair.

In the present study, we created two different sized defects (small and large) at the rotator cuff tendon-bone attachment in the rat and assessed biomechanical properties and structural healing of these defects at different healing time points (no healing and 3- and 8-weeks of healing). We hypothesized that small and large defects would both result in fibrotic scar tissue formation, but that defect size matters; we expected that large defects would propagate into complete tears with evidence of tendon retraction and have diminished healing outcomes compared to small defects (i.e., decreased mechanical properties, bone quality, and organization).

## METHODS

### Surgical Procedure and Gross Morphology

Mature, Long Evans female rats (N = 32 total, n=8 per defect size for *in vivo* healing; n=10 per defect size for mechanical testing post-healing; and n=14 per defect size for non-healed injury mechanics, former breeders, 299±54g) were used under the accordance of the University of Delaware Institutional Animal Care and Use Committee. Under anesthesia (isoflurane carried by oxygen), rats underwent a bilateral surgical procedure to model small- and large full-thickness tears of the infraspinatus tendon-bone attachment using 0.3mm (small, ∼33% damage) and 0.75mm (large, ∼66% damage) punch biopsies (Robbins, Chatham, NJ USA).^18,19^ A 2-cm incision was made craniolateral to the glenohumeral joint and was followed by a horizontal incision to detach the deltoid from the acromion. A suture was passed under the acromion to elevate the scapula and expose the rotator cuff tendon-bone attachments. The infraspinatus tendon-bone attachment was located following internal shoulder rotation, the joint was stabilized, and the defect was created. Small or large punch biopsies were used at the center of the tendon-bone attachment to create full-thickness defects, which permeated and removed the fibrocartilaginous attachment and underlying cortical bone. The shoulder (left or right) that received a small or large defect was alternated between rats using controlled randomization. The muscle layer and skin were closed using simple interrupted resorbable sutures (5-0 Vicryl, Ethicon Inc., Somerville, New Jersey). Rats were given subcutaneous buprenorphine (0.05mg/kg) as analgesia and local bupivacaine hydrochloride (0.05mg/kg) as an anesthetic pre-operatively and postoperatively, respectively. Animals were closely monitored during recovery and twice daily for three days post-operatively. Rats were housed in cages (2 rats maximum per cage) with enrichment and unlimited access to food and water. Rats were euthanized via carbon dioxide asphyxiation and thoracotomy at 3- or 8-weeks post-surgery and entire shoulder complexes were dissected to visualize the infraspinatus attachment. Gross morphology images were taken using a digital camera (DSLR, Nikon). The 3-week time point was used to represent the inflammatory stage of the injury, and the 8-week time point was used to represent the remodeling stage (Figure 1, study design).

**Figure 1.**
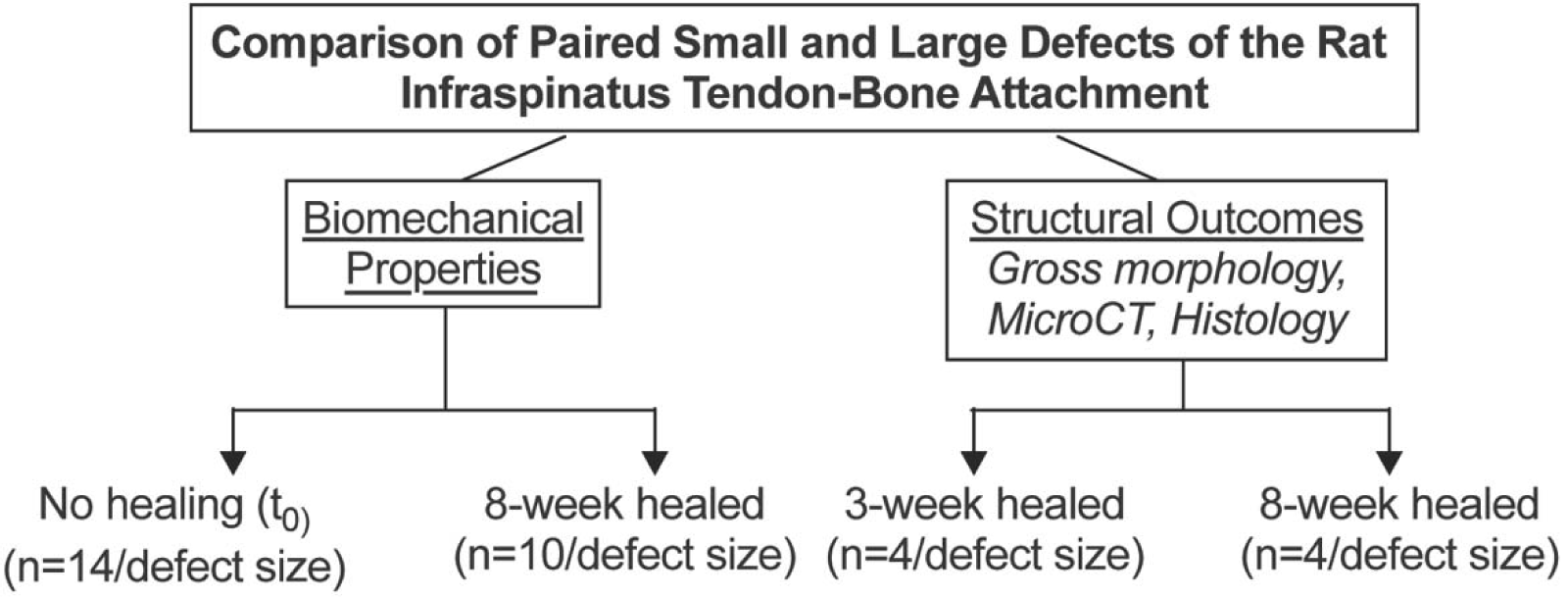
Study design. We compared the biomechanical properties and structural outcomes of paired small and large defects in the infraspinatus tendon-bone attachment of Long Evans rats *in vivo*. Biomechanical properties of small and large defects were assessed with no healing (t_0_) and after 8-weeks of healing. The structural outcomes from gross morphology, micro-computed tomography (microCT), and histology were assessed after 3-weeks and 8-weeks of healing.

### Biomechanics

Mechanical testing was performed to compare mechanical outcomes associated with two different defect sizes both with no healing (t_0_, n=14 per defect size) and following a duration of healing (8-week time point, n=10 animals). A non-healed injury was mimicked in euthanized rats that were stored frozen until time of mechanical testing. Stereoscopic images were taken of the lateral and inferior sides of the attachment (DSLR, Nikon) and ImageJ was used to measure the width and thickness of these sides, respectively. The remaining intact attachment width was calculated by subtracting the defect diameter from the total attachment width for non-healed injuries. For non-healed injuries, the attachment was assumed to be rectangular to approximate cross-sectional area (CSA). For healed attachments, the CSA was assumed circular and measured from the lateral attachment width because there was a large circular scar around the healed attachments rather than rectangular “borders” that were more easily visualized in the non-healed injuries. The humeral head was secured using steel wire (32AWG) to prevent growth plate failures. Distal humeri were potted in 1.5mL centrifuge tubes using polymethyl methacrylate (Ortho-Jet BCA, Lang Dental) or DP100 epoxy (3M, Scotch-Weld). Infraspinatus muscles were removed, and tendons were affixed in thin-film grips lined with PBS-soaked tissue paper (Kimwipes, Kimberly-Clark). Tendons were positioned in a materials test stand at 0-degree shoulder abduction. A 0.1N tare load was applied, and gauge length was recorded before uniaxial tensile tests began (Instron 5943, Norwood, MA). The test protocol was as follows: 10 preconditioning cycles from 0.025-0.075mm at a rate of 0.01mm/sec, followed by 90 seconds of rest, and ramp to failure at 0.01mm/sec. Ultimate load, stiffness, ultimate stress, strain at ultimate stress, Young’s modulus, area under the curve (AUC) at ultimate stress, and AUC of the last five load-displacement preconditioning cycles (energy loss) were computed (MATLAB, Natick, MA).^18^

### Microcomputed Tomography

Immediately following euthanasia and dissection, shoulders allocated for histology and micro-computed tomography (micro-CT) from 3- and 8-week time points (N = 8 rats total, 4 shoulders per defect size per time point) were fixed in 4% paraformaldehyde for 24-48 hours then scanned in air using micro-CT (1276 Skyscan; Bruker, 55kV source voltage, 200μA source current, 10.6µm image pixel size, 1200ms exposure). MicroCT images were reconstructed and aligned with DataViewer (Bruker). Bone morphometric properties for the defect area, greater tubercle, and proximal epiphysis were measured using Dragonfly software (Object Research Systems Inc., Figure 4A and S1). The defect area was defined following alignment of the X- and Y-planes for a normal view of the defect in the Z-planar view. Using the 2D painter tool in Dragonfly, a circle of diameter equal to the diameter of the punch biopsy was digitally painted on the first slice where the full defect was observed in the z-plane. A cylinder-shaped region of interest (ROI) was then filled into the defect using a consistent cylinder height (370µm). Split and Otsu functions were used to separate mineral (bone) and non-mineral within this cylindrical ROI. Bone volume (BV) was divided by the total volume (TV) of the cylinder to calculate (BV/TV) ratio. Epiphyseal ROIs (total epiphysis and greater tubercle) were isolated using consistent anatomical landmarks, and the bone analysis function was used to fill holes smaller than 300μm in order to obtain the TV of the epiphysis. Trabecular and cortical bone were separated using the Kohler method (maximum trabecular thickness=150μm).^35^ Total, trabecular, and cortical bone ROIs were isolated and analyzed using the Bone Analysis tool in Dragonfly (Figure 4A).

**Figure 2.**
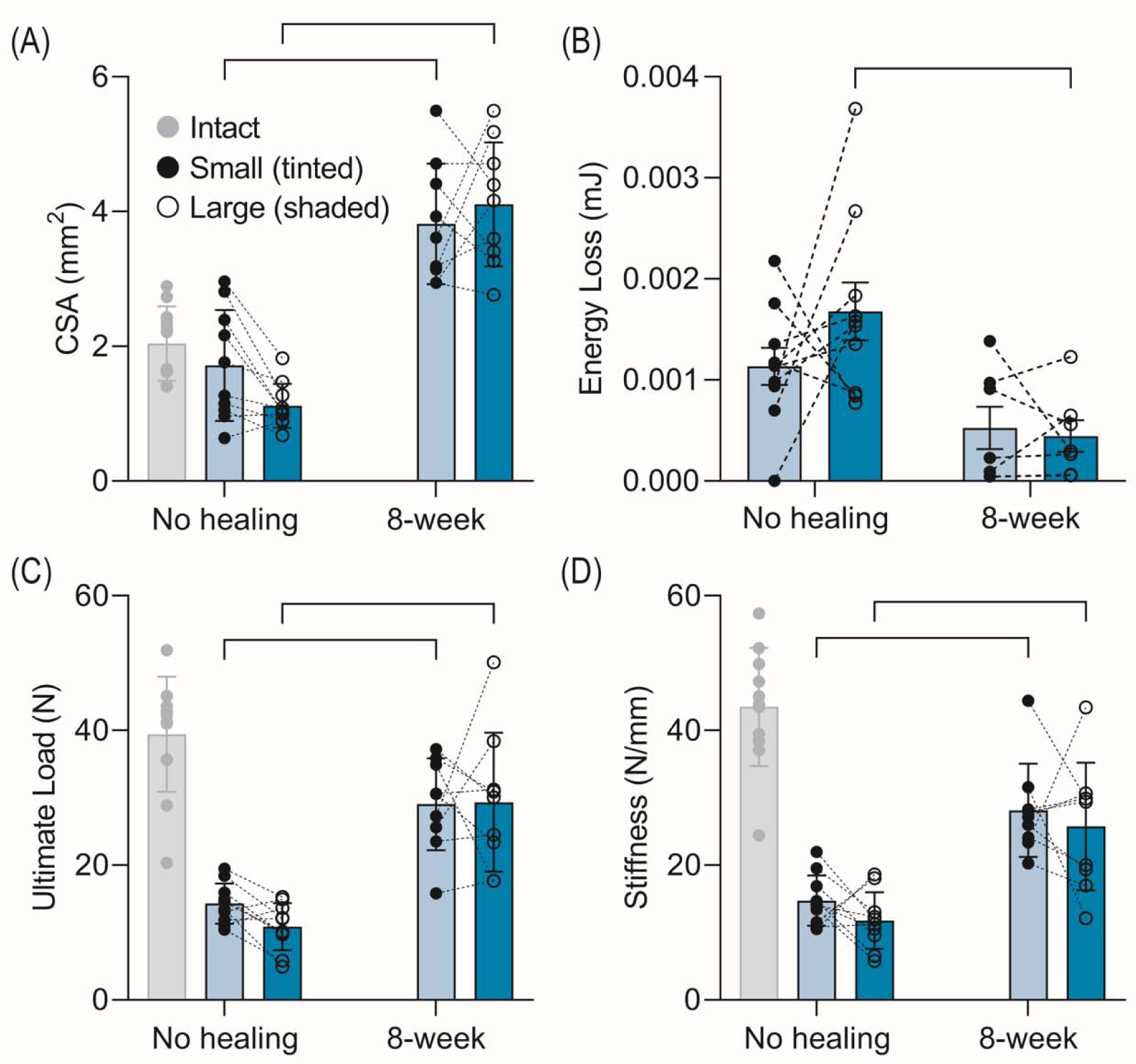
Critical defects that do not regenerate the intact mechanical properties of the tendon-bone attachment may be smaller than one-third of the attachment. (A) Cross sectional area (CSA), (B) energy loss, (C) ultimate load, and (D) stiffness for non-healed and healed small and large defects. Gray data: Intact attachment mechanical properties from Locke et al.^18^ Bars: significant difference (p<0.05). Dashed lines: paired data points. Data are presented as mean ± standard deviation.

**Figure 3.**
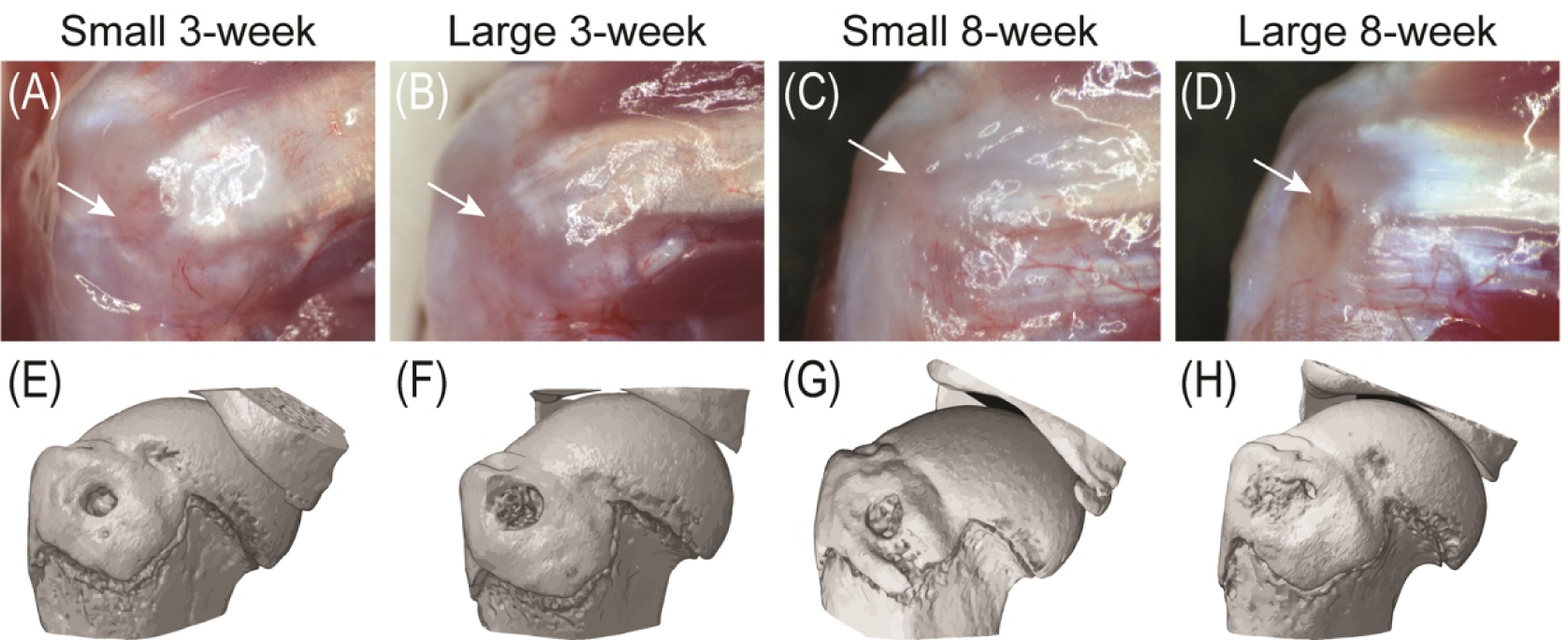
Increased neovascularization, fibrosis, and cortical bone loss were observed for large defects compared to small defects. (A-D) Gross morphology of the defect sites (white arrows) at the time of dissection for small and large defects. (E-F) Representative micro-computed tomography (microCT) 3-dimensional reconstructions of the defect site at the greater tubercle of the humeral head for small and large defects following 3- and 8-weeks of healing.

**Figure 4.**
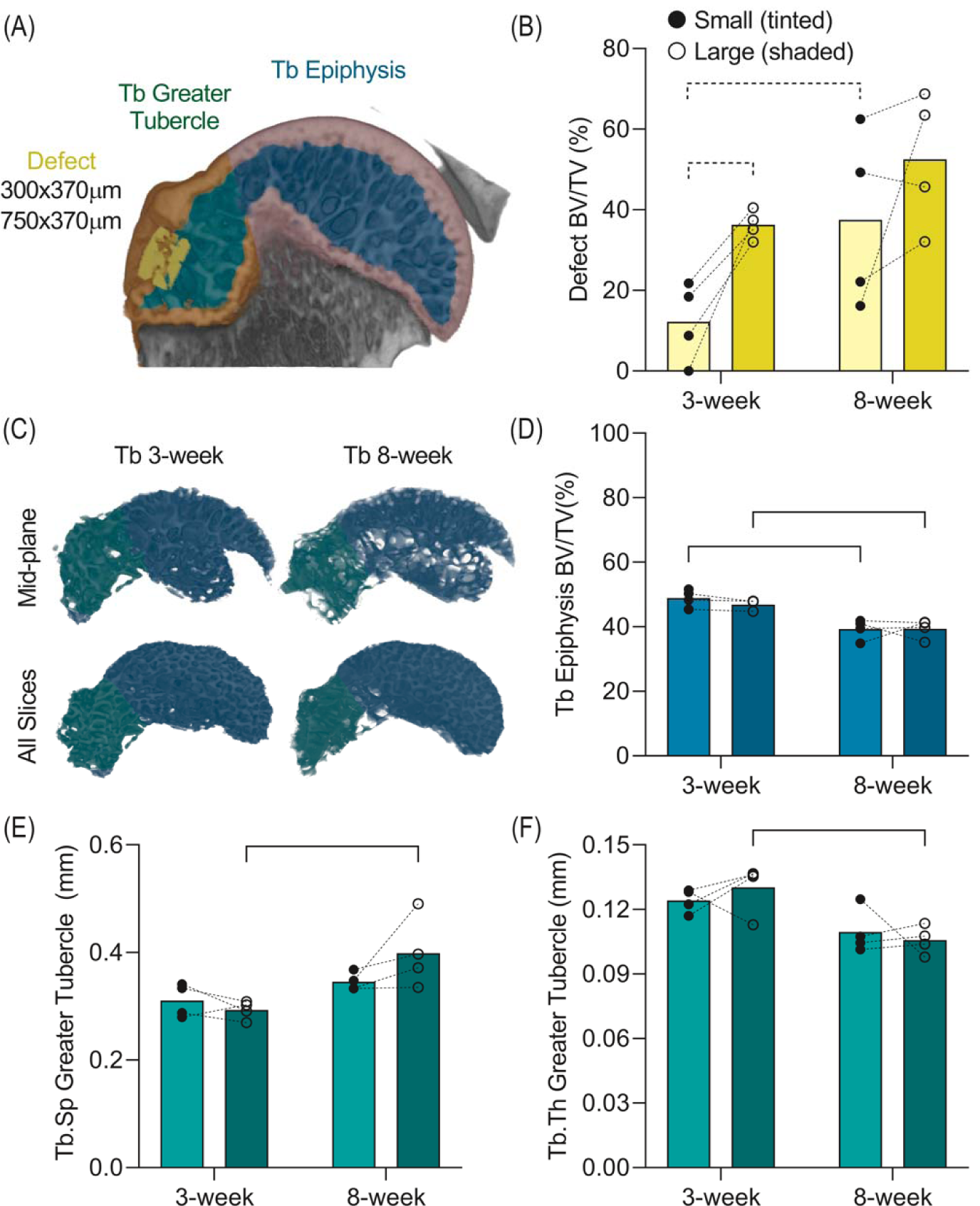
Bone properties increased locally at the defect site and decreased away from the defect site from 3-weeks to 8-weeks. (A) Representative microCT 3-dimensional (3D) reconstruction showing the regions of interest (ROIs) for trabecular (Tb) and cortical (Ct) bone segmentation and analysis: Tb epiphysis (blue), Ct epiphysis (purple), Tb greater tubercle (green), Ct greater tubercle (orange), and defect area (yellow). The epiphyseal ROIs (Tb and Ct) included data from all ROIs. (B) Defect area bone volume (BV) to tissue volume (TV) ratio (BV/TV) was increased in large compared to small defects overall. (C) Representative microCT 3D reconstruction at the mid-frontal plane and of the total epiphyseal Tb bone, showing reduced bone quality from 3- to 8-weeks. (D) Tb epiphysis BV/TV, (E) Tb separation (Sp) and (F) Tb thickness (Th) at the greater tubercle. Dashed bars: trend (p<0.1). Bars: significant difference (p<0.05). Dashed lines: paired data points. Data presented in bar charts are means.

### Histology

After microCT, samples were decalcified in Ethylenediaminetetraacetic acid (EDTA), paraffin embedded, and sectioned at 6µm thickness. Sections were stained with nuclear stain 4’,6-diamidino-2-phenylindole (DAPI) and imaged at 3 individual locations within the attachment (0.024mm^2^/image) to quantify cell number, aspect ratio (nAR), minor axis, major axis, equivalent diameter, perimeter, solidity, eccentricity, orientation and area of nuclei using a custom MATLAB code (Figure 5A-C).^36,37^ Image replicates (3 per sample) were averaged per sample then used for statistical comparisons between defect groups. Hematoxylin & Eosin (H&E) was used to assess fibrosis, cellularity, and vascularity, and Picrosirius Red was used to assess collagen organization.

**Figure 5.**
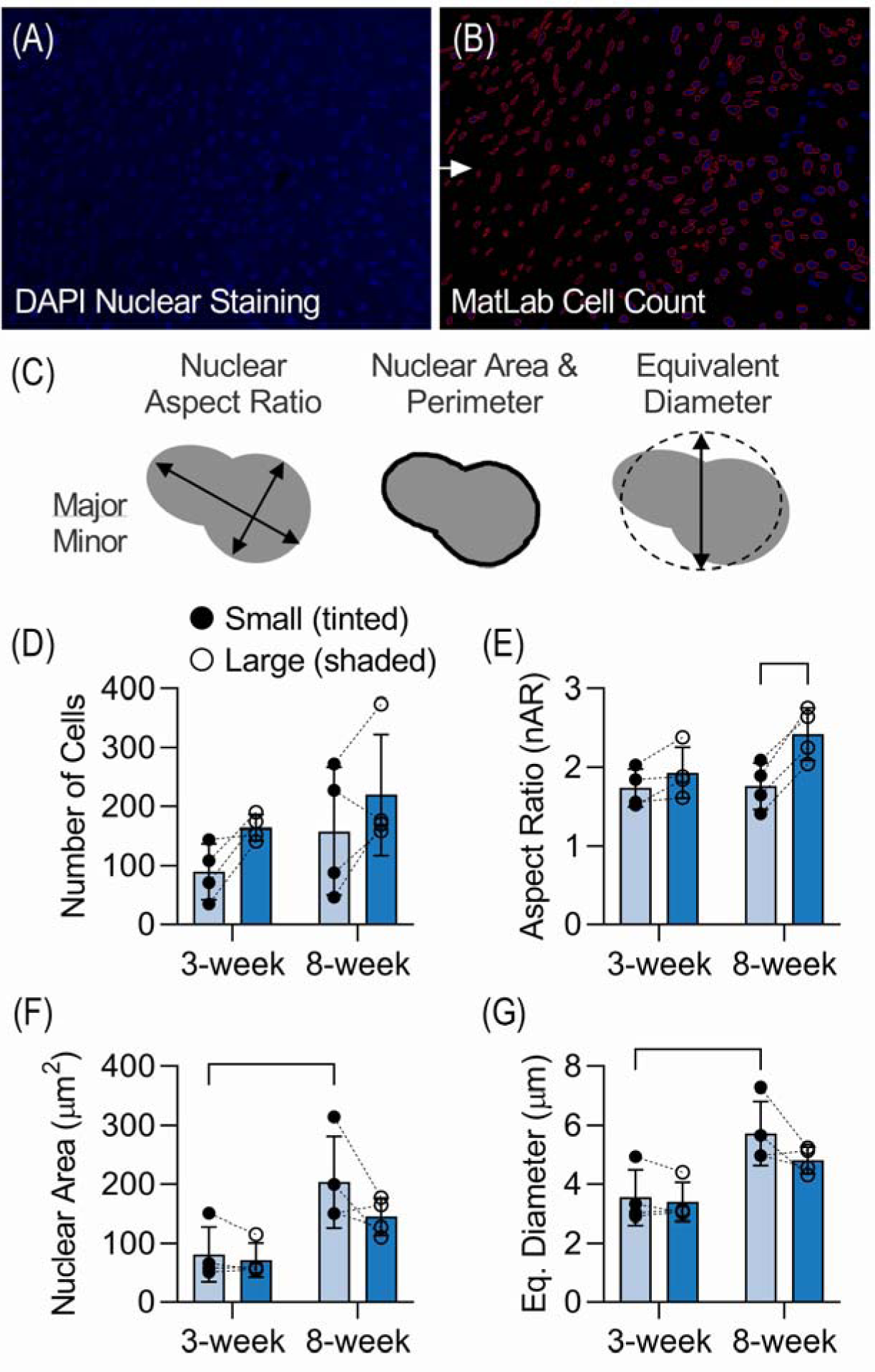
Cellularity and nuclear morphology were moderately impacted by defect size. (A) DAPI stained sections from within the injured attachment were imaged at 40x. Three technical replicates per biological sample were imaged and (B) a custom Matlab code was used to digitize and auto-segment the nuclei. The code was used to calculate and average (C) nuclear properties for each technical and biological replicate. (D) Cellularity, (E) nuclear aspect ratio, (F) nuclear area, and (G) equivalent (Eq) diameter were measured and compared between defect sizes and healing durations. Bars: significant difference (p<0.05). Dashed lines: paired data points. Data are presented as mean ± standard deviation.

### Statistical Analyses

Two-way ANOVA with repeated measures and Sidak’s multiple comparison tests were used to compare mechanical outcomes, bone morphometry from microCT, and cell nuclear shape of paired small and large defects at t_0_ (mechanics only), 3- or 8-weeks of healing. All statistics were performed using Prism (version 8, GraphPad). Mechanical properties of intact attachments from Locke et al. were used for qualitative, not statistical, comparisons.^18^

## RESULTS

All animals tolerated bilateral surgery well.

### Mechanical Properties Increased with Healing

Four non-healed rats and one healed rat were excluded due to errors during data acquisition, dissection, and abnormal defect placement; therefore, n=10 per defect size were analyzed in the non-healed group, and n=9 per defect size were analyzed for the 8-week healed group. Two additional rats in the healed group were excluded from energy loss comparisons, but not load to failure comparisons, due to data acquisition errors during preconditioning.

Attachment CSA decreased as non-healed defect size increased, and following 8-weeks of healing, CSA was significantly increased for both defect groups (Figure 2A). In the non-healed group, ultimate load and stiffness were reduced and energy loss was higher but not significantly different in large compared to small defects (Figure 2B-D). Ultimate load and stiffness were reduced in both non-healed defect groups compared to intact attachments (Figure 2C-D), corroborating previous findings from our group.^18,19^

After 8-weeks of healing, both small and large defects had increased ultimate load and stiffness compared to non-healed defects; however, healed defects did not regain comparable properties to native, uninjured attachments (Figure 2C-D). Energy loss was reduced for healed defects compared to defects with no healing (Figure 2B). No differences were observed for Young’s modulus, ultimate stress, AUC at ultimate stress, or strain at ultimate stress between defect sizes or healing duration (Table S1).

### Gross Morphology and Bone Properties Were Affected by Defect Size and Healing Duration

From gross morphology, no tear propagation into complete tears or tendon retraction was observed for either defect size (Figure 3A-D), although increased neovascularization and fibrosis at the defect site were found for large compared to small defects (Figure 3A-D, white arrows). Large defects had increased fibrosis at 8-weeks compared to 3-weeks (Figure 3A-D).

Additionally, we qualitatively observed less cortical bone for large defects compared to small defects at 3-weeks (Figure 3E-F).

The amount of bone within the defect increased over time for both defect sizes (Fig 4) and bone volume fraction (BV/TV) was higher in large compared to small defects especially at 3-week (Figure 4B and S1, p=0.02). Conversely, trabecular morphometry of the humeral head (trabecular epiphysis, Figure 4C-D) and of the greater tubercle (Figure S2) decreased with increased healing time but did not vary between defect sizes. In the greater tubercle, trabecular separation (Figure 4E, p < 0.01) increased and trabecular thickness (Figure 4F, p<0.01) decreased from 3-to 8-weeks but did not differ between defect sizes.

### Tissue Cellularity and Structure Were Mildly Different Between Defect Groups

Within the defect, cellularity was increased for large defects compared to small defects overall (Figure 5D, p=0.02). Large defects had significantly increased nuclear aspect ratio (Figure 5E) and eccentricity (Table S2) compared to small defects overall and at 8-weeks of healing (Figure 5E). The nuclear area (Figure 5F), equivalent diameter (Figure 5G), major axis, minor axis, and perimeter (Table S2) were all increased from 3- to 8-weeks of healing overall and in small defects from 3- to 8-weeks. The nuclear solidity and orientation were not affected by defect size or healing time (Table S2).

Both defect sizes had evidence of vascularity within the injured attachment (Figure 6A-B) and more vessels were observed for large defects compared to small defects at 3-weeks of healing (Figure 6B). Additionally, at 3-weeks, both defects had rounded nuclei within the injury region rather than elongated tendon fibroblasts, indicating the presence of macrophages or other cell populations (Figure 6A-B). From 3- to 8-weeks, both small and large defects had evidence of fibrosis surrounding the defect region and reduced appearance of vessels (Figure 6C-D). Qualitatively, we observed that small and large defects had mixed collagen organization after 3-weeks of healing (Figure 6E-F), while large defects resulted in more unorganized collagen compared to small defects after 8-weeks of healing (Figure 6G-H).

**Figure 6.**
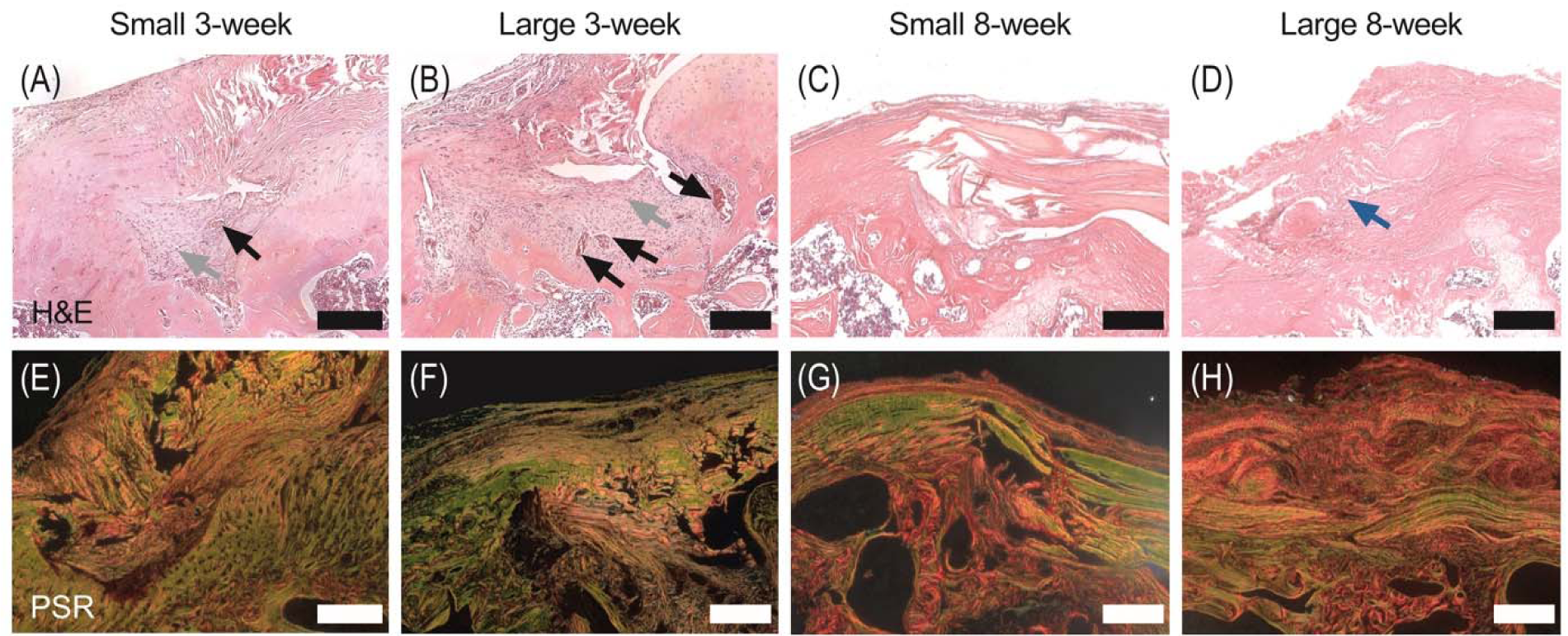
Large defects had increased vasculature and scar tissue at 3-weeks and 8-weeks of healing, respectively. (A-D) Hematoxylin and Eosin (H&E) and (E-H) Picrosirius Red (PSR) were used to assess tissue micro-structure and collagen organization. (A-B) At 3-weeks, both small and large attachments had evidence of vascularity (black arrows). Rounded nuclei (gray arrows) were observed at 3-weeks in both sized defects. (C-D) Following 8-weeks of healing, both defects had evidence of a fibrotic scar formation, with more observed in large defects (blue arrow). (E-F) Small and large defects had mixed organization of collagen after 3-weeks of healing. (G-H) Qualitatively, large defects compared to small defects resulted in more unorganized collagen at 8-weeks. Scale bars = 100µm.

## DISCUSSION

In our study, we showed that the size of full-thickness rotator cuff defects did not play a significant role in mechanical outcomes, but did affect the gross morphology, bone morphometry, cellularity, and structure of the tendon-bone attachment. Surprisingly, and counter to our hypothesis, we observed similar mechanical properties of both defect sizes after 8-weeks of healing. Neither defect group regained the native mechanical properties of the healthy attachment, indicating that the attachment does not fully regenerate within 8-weeks following injury. Therefore, one-third or even less than one-third of the width of the tendon-bone attachment may be a critical defect in the rat rotator cuff. It is possible that a longer duration of healing could have led to restored mechanical properties comparable to the intact tendon-bone attachment. Additionally, differences in mechanical properties between defect sizes may have occurred at shorter durations of healing. Although we did not find a mechanical difference between defect sizes, we did observe several spatiotemporal differences in attachment bone properties, cellularity, and structure between defect sizes. We showed that the size of full-thickness rotator cuff defects was in part associated with healing outcomes.

Our model of full-thickness injury is similar to crescent-shaped tears that commonly present in the clinic and lead to tear propagation.^1,3^ Tear propagation has been correlated with increased strain at the defect site.^20^ In our injury model, although we observed multiple decreased mechanical properties between non-healed small and large defects compared to healthy attachments and have previously observed strain concentrations at the defect site^18^, we did not observe tear propagation in either defect size following 8-weeks of healing *in vivo* under normal physiologic loading. Increased mechanical loading at the attachment to the point of overuse, however, may increase the likelihood of tear propagation even for small defects.^38,39^ In our study, we did not challenge the injured attachment beyond the initiation of a defect. Physiologic loading in this study may have resulted in a protective healing response that prevented tear propagation into a complete tear and supported a similar increase in mechanical properties of both defects during healing.^40^ Future studies could investigate the role of acute or repetitive loading on attachment damage and tear propagation by modulating muscle loading and attachment mechanobiology (e.g., treadmill running or muscle stimulation).^41^

Although we observed that the duration of injury affected healing outcomes more so than defect size, defect size lead to several differences in gross morphology, bone properties, cellularity, and attachment structure. Rotator cuff tears commonly lead to fibrosis and neovascularization,^19,24^ and, in this study, we similarly observed a qualitative increase in fibrotic tissue and vessel formation with increased defect size. Additionally, various dynamic cell behaviors, such as migration, proliferation, and differentiation, as well as abnormal cell phenotypes, such as rounded cells, have been observed following tendon injury.^42,43^ Notably, we observed a dynamic increase in cell number and structural characteristic of the attachment, including fibrosis and collagen organization, that were dependent both on defect size and healing time, concurring with Lemmon et al. but contrasting other injury models.^19,30,31^ A decrease in cell number and increased collagen organization may be observed with further healing duration, yet comparable tendon remodeling has been observed between 8- and 12-weeks,^44^ suggesting that 8-weeks is an appropriate time point to assess attachment remodeling. These findings suggest that the rat attachment may adapt to the increased tear size in attempt to remodel the larger injury.

Nuclear aspect ratio (nAR) has been used previously as read out of mechanical force transmission experienced by cells, and changes in nuclear shape can trigger epigenetic changes and influence gene expression.^41,42,45,46^ Previous studies have shown that tendon injury leads to an acute decrease in nAR compared to healthy tendons and that nAR increases with increased healing duration, indicating a mechanobiological response to injury and healing.^42^ Interestingly, in this study, we observed changes in nAR of local fibroblasts that were dependent on injury size. Whether large defects result in increased tissue-scale strain compared to small defects and if the mechanical differences between defect sizes lead to differences in nAR remain unknown.^18^ Alternatively, increased nAR, as well as nuclear area and equivalent diameter, in larger defects may suggest increased differentiation of local or extrinsic stem cells into fibroblasts.^42,47^ Ultimately, we suggest that the dynamic nuclear changes observed in this study possibly developed from increased tissue-scale strain with increased defect size and/or altered mechanobiological force transmission with increased tissue remodeling. A further understanding of attachment mechanobiology may help to clarify these cellular adaptations to attachment defect size *in vivo*.^41,48^

### Limitations

The use of this bilateral model allowed us to minimize the potential for abnormal loading on healing outcomes. However, a limitation to this study is that we did not quantify limb preference during gait. Our model was beneficial as we utilized a bilateral defect which allowed us to pair our comparisons using biologically identical shoulders. Additionally, while decreased bone quality following acute tendon injury is common^25^, the nature of our model likely contributed to a decrease in bone volume as the punch biopsy permeated through cortical bone.^18,19^ Lastly, the use of an animal model to study rotator cuff tears has limitations when translating to humans due to differences in joint loading and healing. Although rats are a commonly used rotator cuff model, the use of rats in our study could potentially be a limiting factor. Rats have an increased propensity to heal with limited scar formation and are also quadrupeds that weight-bear on their forearms unlike humans. Although they perform overhead reaching and grasping, this is not identical to human biped motion.

It is important to understand how defect size can influence the mechanical and structural outcomes of attachment healing. The use of animal models to study the mechanical consequences and healing outcomes of critical defect size(s) can improve our preclinical understanding of early rotator cuff disease. This defect model can serve as a tool to understand the native healing process of full-thickness tears and as a testbed for preclinical testing of biomaterials that could be used to mitigate or delay surgical intervention of early rotator cuff disease.

## ACKNOWLEDGEMENTS

Funding: Delaware Space Grant Consortium (NNX15AI19H), the UD Dare to be FIRST Research Experience for Undergraduates, the National Science Foundation Grant (1460757), the University of Delaware Doctoral Fellowship, the Eunice Kennedy Shriver National Institute Of Child Health & Human Development of the National Institutes of Health (K12HD073945), and the National Institute of General Medical Sciences of the National Institutes of Health (P30GM103333). We would like to thank Francis Karani for animal handling and Dr. Gwen Talham for veterinary care.

## Notes

### Competing Interest Statement

The authors have declared no competing interest.

### Summary of Updates

We have created a model of acute, full-thickness tears of varying width (i.e. small and large). This terminology was clarified and addressed throughout the manuscript.

