## Supplemental data for "Mechanics and Differential Healing Outcomes of Small and Large Defect Injuries of the Tendon-Bone Attachment in the Rat Rotator Cuff"

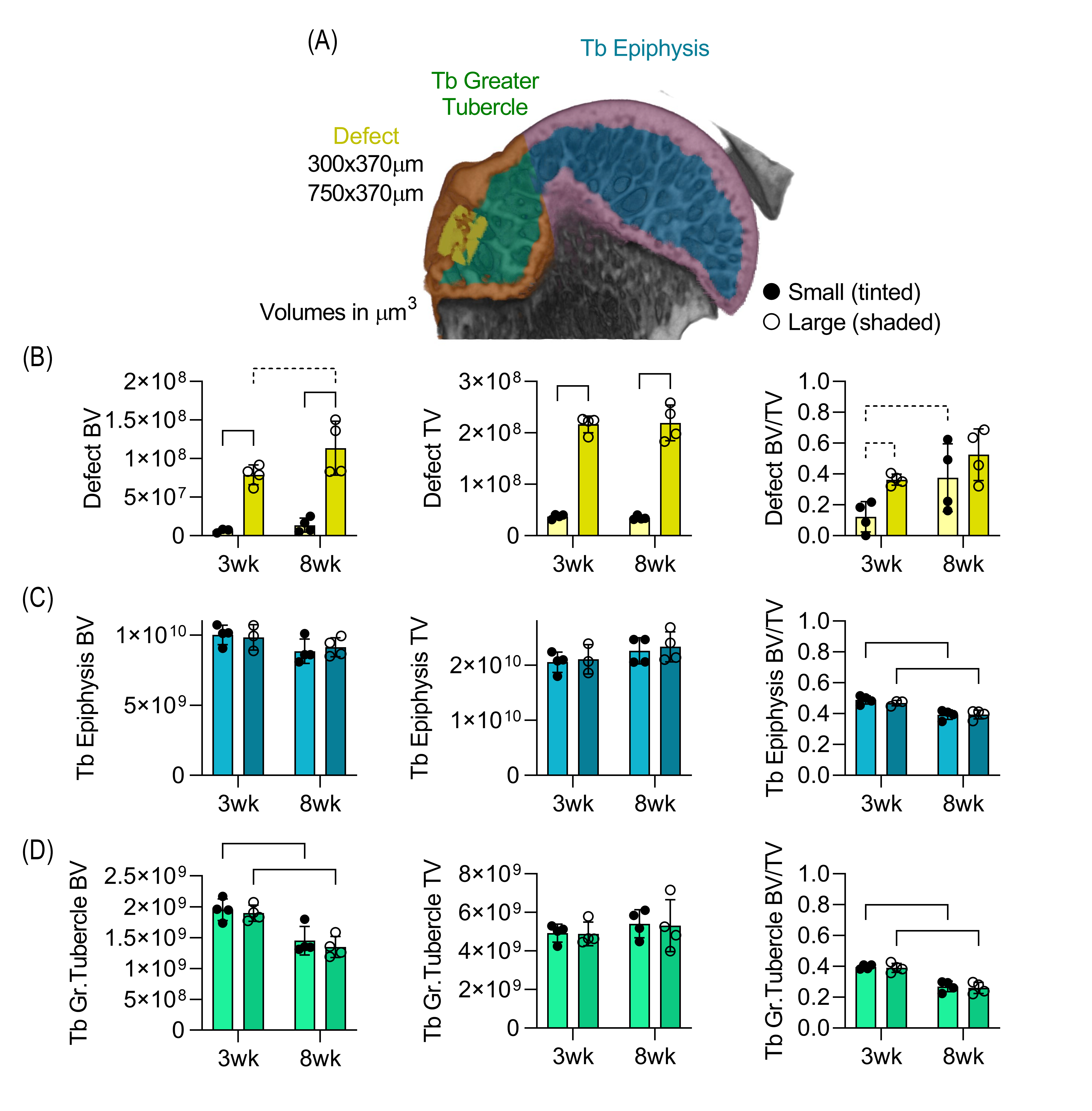
 **Figure S1.** Bone morphometry changes between small and large defects at 3- and 8-weeks of healing. (A) Representative microCT 3-dimensional (3D) reconstruction showing the regions of interest (ROIs) for trabecular (Tb) and cortical (Ct) bone segmentation and analysis: Tb epiphysis (blue), Ct epiphysis (purple), Tb greater tubercle (green), Ct greater tubercle (orange), and defect area (yellow). The epiphyseal ROIs (Tb and Ct) included data from all ROIs. Bone volume (BV), total volume (TV), and BV/TV ratio for each region of interest: (B) defect area, (C) epiphysis Tb, and (D) total greater tubercle Tb. Bars: significant difference (p<0.05). Dashed lines: paired data points. Data are presented as mean.

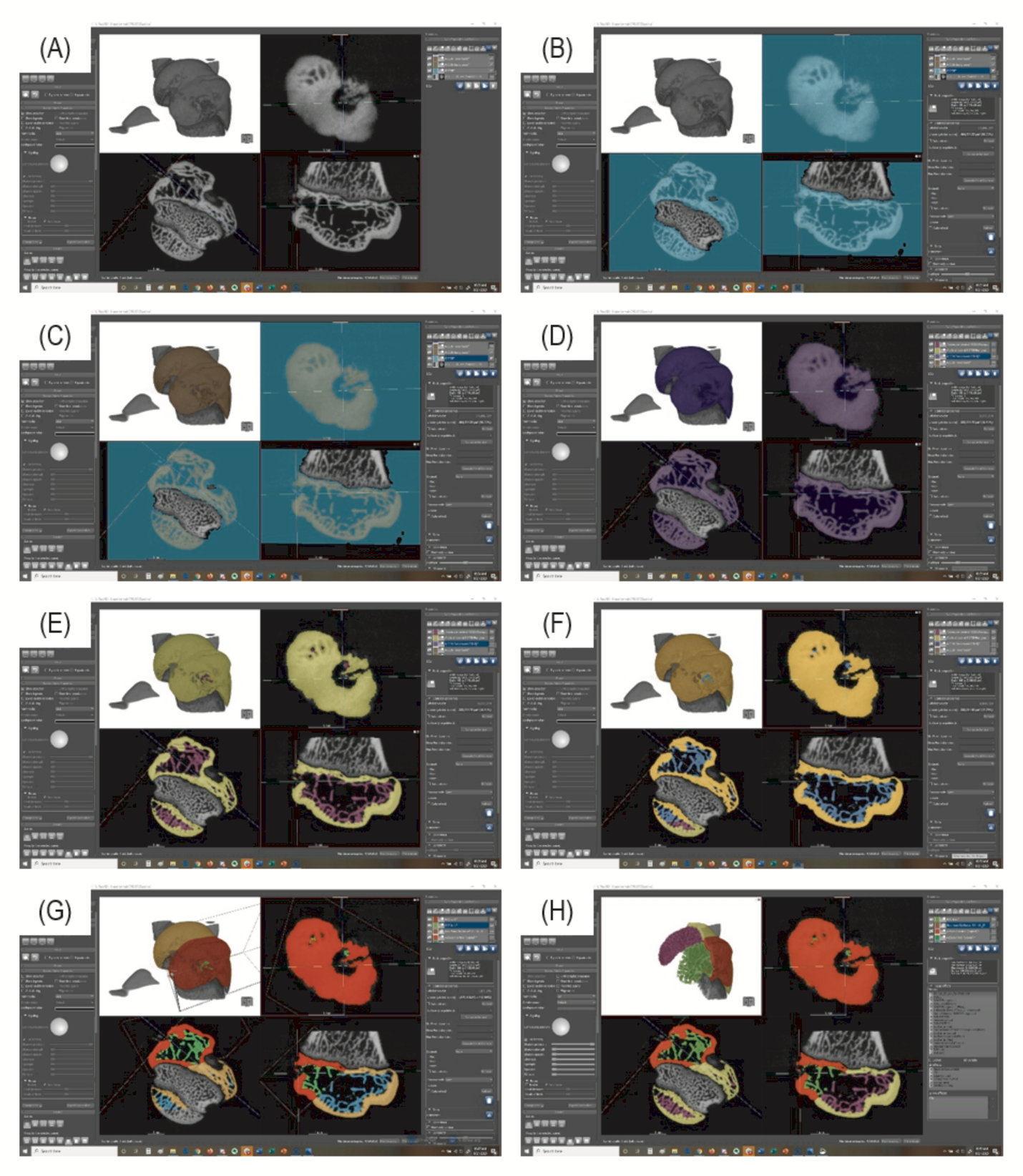

**Figure S2.** Dragonfly was used to distinguish mineral from empty space and separate cortical and trabecular bone. (A) The sample was rendered in a 3-dimensional space. (B) A region of interest (ROI) was created that excluded the metaphysis, diaphysis, scapula, and acromion. (C) Using the “Split at Otsu” function, pixels with light values were separated into a new channel that represented mineral content. (D) Using the bone analysis tool, the total volume was calculated. (E) Bone analysis used an input of 150µm trabecular thickness to separate ROIs for epiphysis cortical bone (yellow) and epiphysis trabecular bone (purple) and were used for epiphysis BV/TV. (F) The greater tubercle was isolated and ROIs were created for cortical bone (orange) and trabecular bone (blue). (G) The anatomical neck was used as a consistent landmark to separate the humeral head and greater tubercle, and then the greater tubercle cortical bone (red) and greater tubercle trabecular bone (green) were separated. The greater tubercle ROIs were then used for the final step of bone analysis. (H) Epiphysis and greater tubercle ROIs could then be visualized through a clip box.

**Table S1.** No differences were observed for Young’s modulus, ultimate stress, AUC at ultimate stress, or strain at ultimate stress from 0- to 8-weeks of healing time (t) or between small (S) and large (L) defect sizes. X: interaction between time and size. Data are presented as mean ± standard deviation.

| **t** | **Size** | **Young's Modulus (MPa)** | **Ultimate Stress (MPa)** | **AUC at ultimate stress (J/mm^3^)** | **Strain at ultimate stress (mm/mm)** |
| --- | --- | --- | --- | --- | --- |
| t_0_ | S | 33.82±20.49 | 10.63±6.12 | 2.27±1.34 | 0.41±0.13 |
|  | L | 37.97±11.50 | 9.75±15.69 | 1.58±0.83 | 0.35±0.19 |
| 8wk | S | 31.39±13.77 | 7.94±2.51 | 1.36±0.75 | 0.40±0.12 |
|  | L | 23.72±11.50 | 7.65±3.30 | 1.69±0.83 | 0.51±0.18 |
| t |  |  |  |  |  |
| Size |  |  |  |  |  |
| X |  |  |  |  |  |

**Table S2.** Nuclear morphology outcomes at 3-weeks and 8-weeks of healing time (t) for small (S) and large (L) defect sizes. Single asterisk represents significant differences between time points for specific defect sizes, and two asterisk represents significant differences for time or size overall. X: interaction between time and size. Data are presented as mean ± standard deviation.

| **t** | **Size** | **Major Axis (µm)** | **Minor Axis (µm)** | **Perimeter (µm)** | **Solidity (%)** | **Eccentricity (%)** | **Orientation (°)** |
| --- | --- | --- | --- | --- | --- | --- | --- |
| 3wk | S | 4.75±1.53 | 2.86±0.65 | 12.20±3.62 | 0.94±0.01 | 0.71±0.06 | 3.38±7.64 |
|  | L | 4.84±1.36 | 2.64±0.36 | 12.11±2.93 | 0.93±0.01 | 0.75±0.06 | -7.24±15.75 |
| 8wk | S | 7.63±1.02* | 4.67±1.09* | 20.39±3.04* | 0.93±0.03 | 0.72±0.07 | 4.17±17.00 |
|  | L | 7.65±0.38 | 3.46±0.52 | 18.89±2.00 | 0.91±0.02 | 0.83±0.04* | -3.80±26.60 |
| t |  | ** | ** | ** |  |  |  |
| Size |  |  |  |  |  | ** |  |
| X |  |  |  |  |  |  |  |
